# Adolescent ethanol drinking promotes hyperalgesia, neuroinflammation and serotonergic deficits in mice that persist into adulthood

**DOI:** 10.1101/2021.11.29.469930

**Authors:** Kanza M. Khan, Gabrielle Bierlein-De La Rosa, Natalie Biggerstaff, Selvakumar Govindhasamy Pushpavathi, Suzanne Mason, Michael E. Dailey, Catherine A. Marcinkiewcz

## Abstract

Adolescent alcohol use can permanently alter brain function and lead to poor health outcomes in adulthood. Emerging evidence suggests that alcohol use predispose to pain disorders or exacerbate existing pain conditions, but the neural mechanisms are currently unknown. Here we report that mice exposed to adolescent intermittent access to ethanol (AIE) exhibit increased pain sensitivity and depressive-like behaviors that persist after alcohol cessation and are accompanied by elevated CD68 expression in microglia and reduced numbers of serotonin (5-HT)-expressing neurons in the dorsal raphe nucleus (DRN). 5-HT expression was also reduced in the thalamus, anterior cingulate cortex (ACC) and amygdala as well as the lumbar dorsal horn of the spinal cord. We then found that chronic minocycline administration after AIE alleviated hyperalgesia and social deficits, while chemogenetic activation of microglia in the DRN of *Cx3cr1-cre-GFP* mice reproduced the effects of AIE on pain and social interaction. Taken together, these results indicate that microglial activation in the DRN may be a primary driver of pain and negative affect after AIE.

## 1. Introduction

The adolescent brain is highly sensitive to the effects of ethanol which may predispose individuals to wide range of neuropsychiatric comorbidities including chronic pain, a condition that afflicts approximately 20% of adults in the U.S. (Broadwater and Spear, 2013; Dahlhamer et al., 2018; Nasrallah et al., 2011; Risher et al., 2015; Schindler et al., 2014; Wolstenholme et al., 2017). Comorbidity between ethanol dependence and pain is often attributed to self-medication of symptoms (Brennan et al., 2005; Thompson et al., 2017), but emerging evidence suggests that heavy alcohol drinking and other chronic stressors may exacerbate or even precipitate pain symptoms (Caniglia et al., 2020; de Oliveira et al., 2017; Egli et al., 2012; Fu et al., 2015; Marcinkiewcz et al., 2009). Several recent studies have reported microglial activation and elevated levels of pro-inflammatory cytokines in the brain following adolescent alcohol exposure, all of which may adversely impact synaptic function in central pain circuits. (Bajo et al., 2015; Crews et al., 2017, 2016; Crews and Vetreno, 2014; Cruz et al., 2017; Marshall et al., 2020; Walter and Crews, 2017; Zhao et al., 2013). Together, these converging lines of evidence suggest that microglia may be a common cellular substrate linking alcohol use to chronic pain disorders (Grace et al., 2018; Sawicki et al., 2019; Yi et al., 2021).

The dorsal raphe nucleus (DRN) is enriched in serotonin (5-hydroxytryptamine; 5-HT) neurons that play a critical role in the regulation of pain and affect (Akil and Mayer, 1972; Dugué et al., 2014; Griffith and Gatipon, 1981; Horie et al., 1991; Mayer and Liebeskind, 1974; Oliveras et al., 1979). Pro-inflammatory stimuli can precipitate 5-HT neuronal degeneration, induce cfos expression, and alter 5-HT neuronal activity in the DRN (Brambilla et al., 2007; Hochstrasser et al., 2011a; Hollis et al., 2006; Manfridi et al., 2003), suggesting that chronic alcohol may promote hyperalgesia and negative affect by activating neuro-immune mechanisms in the DRN that ultimately perturb 5-HT neurotransmission. In the present study, we examined the effects of adolescent intermittent access to ethanol (AIE) on pain and affective behaviors, 5-HT expression, and neuroinflammation in the DRN and other brain regions that have been implicated in nociception. Persistent hyperalgesia was accompanied by microglial activation in the DRN and median raphe nucleus (MRN) and reduced 5-HT expression in the DRN. We also observed a significant reduction in 5-HT expression in pain circuits that receive 5-HT input from the DRN including the thalamus, anterior cingulate cortex (ACC), amygdala and dorsal horn (DH). This AIE-induced hyperalgesia was alleviated by minocycline treatment after alcohol cessation, while chemogenetic activation of microglia in the DRN was sufficient to induce hyperalgesia and social deficits in ethanol-naïve mice. Together, these results suggest that AIE may predispose to chronic pain disorders later in life by activating microglia in the DRN.

## 2. Methods

### 2.1 Animals

All procedures were carried out in accordance with the ethical guidelines for the use of animals in research and approved by the Institutional Animal Care and Use Committee (IACUC) at the University of Iowa. Adolescent male C57BL/6J mice used in voluntary ethanol access experiments had *ad libitum* access to an irradiated diet (Prolab IsoPro 3000) and were maintained on a reverse light-dark cycle (lights off at 07:30). Adult male *Cx3cr1-creER* mice (Jackson Laboratories, Stock #021160) were used in DREADD experiments and maintained on a standard light-dark cycle (lights on at 06:00) with *ad libitum* access to standard chow. Mice were randomly assigned to a treatment group and data was analyzed by trained personnel blinded to the treatment.

### 2.2. Drugs

Ethanol solutions (20% w/v) were prepared in tap water from 95% ethyl alcohol (Decon Labs). Minocycline HCl (Sigma) was dissolved in distilled water (0.0416mg/mL) and administered via drinking bottles at 3 hours into the dark cycle for an average daily dose of 10 mg/kg. Tamoxifen (TM; Sigma) was dissolved in Sunflowerseed Oil and administered at a dose of 75 mg/kg (i.p.). Clozapine N-oxide (CNO) dihydrochloride (Hello Bio) was dissolved in sterile saline and administered to mice by intraperitoneal injection (3 mg/kg, i.p.) 45 min before behavioral testing.

### 2.3. Adolescent intermittent access to ethanol (AIE)

Mice were exposed to an intermittent access to ethanol paradigm as previously described (Hwa et al., 2011). Briefly, mice were individually housed and presented with two 50mL plastic tubes that were fitted with No. 5 rubber stoppers. Three hours into the dark cycle on Mondays, Wednesdays, and Fridays, mice were presented with tubes containing ethanol or water for 24 hr for 4 weeks starting at postnatal day (PND) 25.

Placement of the ethanol and water bottles was alternated daily to control for side preferences. Ethanol intake (g/kg/24h) and preference were calculated after each drinking session for a daily average of 14.04 ± 1.15 g/kg/day.

### 2.4. Stereotaxic surgeries

All surgeries were performed using aseptic technique. Before the surgery, mice were deeply anesthetized with 3% isoflurane and placed on a heating pad in a stereotactic frame. During surgery, isoflurane was maintained at 1.5-2%. To selectively target microglia, AAV encoding Cre-inducible Gq-coupled DREADDs (AAVDJ-ef1α-DIO-hM3Dq-mCherry, Stanford viral vector core) were microinjected into the DRN of *Cx3cr1-creER* mice using a 1 µL Neuros Hamilton syringe at a rate of 100 nl/min. After infusion, the needle was left in place for an additional 10 minutes to allow diffusion in the target area. Coordinates from bregma were AP: -4.65, ML: 0, DV: -3.3, at an angle 23.58° to ML. The incision site was closed, and mice were returned to their regular housing room.

### 2.5. Behavior

#### 2.5.1. Von Frey

Mice were placed inside a clear acrylic box (100 × 100 × 150 mm) and acclimatized to the test apparatus which consisted of a raised wire mesh platform (holes in mesh were 5×5 mm) for at least 3-h on two consecutive days (Warwick et al., 2019). Mechanical sensitivity was evaluated by applying von Frey filaments (Stoelting, Wood Dale, IL) of varying strength to the plantar surface of each hindpaw. The number of responses to each filament out of 5 applications was recorded and used to calculate the 50% threshold.

#### 2.5.2. Hargreaves

Mice were placed in a clear acrylic box (100 × 100 × 150 mm) on a IITC Life Science heated base (Model 400; temperature maintained at 30^0^C) and were acclimated to the test apparatus for 2 consecutive days before testing. Thermal sensitivity was evaluated by focusing a light beam to generate heat on the plantar surface of each hindpaw. The time required for the stimulus to evoke a withdrawal was measured 3 times per paw and averaged to calculate the paw withdrawal latency.

#### 2.5.3. Social interaction

The social interaction test was performed as described previously (Lowery-Gionta et al., 2014; Marcinkiewcz et al., 2014; Moy et al., 2008). Briefly, mice were placed in a clear three-chambered plexiglass arena (∼20 lux) with small openings to allow movement between chambers. Following a 10-min habituation phase, two small metal cages were placed in each side chamber and a novel mouse was placed inside of one of the cages. Placement of the novel mouse was altered between trials to control for side preferences. Behavior was recorded by an overhead camera and scored by a researcher blind to experimental conditions. The time spent interacting with the novel mouse and the empty cage was analyzed over a 10-minute period for each trial.

#### 2.5.4. DREADD experiments

Two weeks following viral infusion into the DRN, *Cx3cr1-creER* mice were injected with tamoxifen (75 mg/kg, i.p.) once daily for 5 days to induce Cre-meditated recombination and DREADD expression in microglia. Mice were used in behavior tests ∼1 month following surgery.

### 2.6. Immunohistochemistry and Microscopy

Mice were anesthetized with Avertin and transcardially perfused with 0.01 M PBS followed by 4% paraformaldehyde (PFA). Brains were extracted, fixed in PFA for 24 h at 4°C, and stored in PBS at 4°C. Brains were sliced on a Leica VT100S at 45μm and stored in 50/50 PBS/Glycerol at -20°C. For each region of interest (ROI), four slices were used across the caudal-rostral axis. Slices were washed in PBS and incubated in 0.5% Triton X-100/PBS for 30 min, blocked in 10% Normal Donkey Serum in 0.1% Triton X-100/PBS and then incubated with the respective primary and secondary antibodies (Supplementary Table 2). Slices were subsequently washed in PBS, mounted on glass slides and coverslipped with Vectashield.

Confocal z-stacks (1 µm) were captured on a confocal microscope (40 sections/z-stack) and converted to maximum projection images using Image J software. Images were analyzed by trained researchers blind to experimental conditions for cell counts, cell body size, and optical density using ImageJ.

### 2.7. Statistical Analyses

Experiments comparing two groups were analyzed with a Student *t-test*, with α < 0.05; social interaction and minocycline experiments were analyzed using a two-way ANOVA and Bonferroni corrections for *post hoc* analyses using Graphpad Prism 8 software. Data are reported as means ± standard error of the mean (SEM).

## 3. Results

### 3.1. AIE promotes hyperalgesia and depressive-like behaviors in adult mice

AIE reduced pain thresholds in the Von Frey and Hargreaves tests at 12 weeks of age, which is indicative of hyperalgesia (Von Frey: t_17_=4.11, p<0.001; Hargreaves t_18_=3.05, p<0.01) (Figure 1B-D). These mice also exhibited significant deficits in social interaction which may be indicative of depressive-like behavior (F_1,17_=12.56, p<0.001, Stranger x Ethanol interaction) (Figure 1E). Surprisingly, there was no effect of AIE on anxiety-like behavior in the elevated plus maze (EPM), although we did observe a decrease in time spent in the center of the open field (t_17_=2.57, p<0.05) which may indicate anxiety-like behavior. There were no differences in locomotor activity in the EPM or the open field test (Supplementary Table 1). There was also no effect of AIE on immobility time in the forced swim test, although we did see an increase in the frequency of inactive bouts (t_18_=2.19, p<0.05) and the average length of inactive bouts (t_11_=2.30, p<0.05 with Welch’s correction) in AIE mice which is consistent with depressive-like behavior (Supplementary Table 1). Together, these results suggest that long-term plasticity in neural circuits mediating pain and affective behavior may occur following AIE. We then examined inflammatory markers in central pain circuits including the DRN and MRN, medullary raphe, thalamus, ACC, hypothalamus, and amygdala.

**Figure 1:**
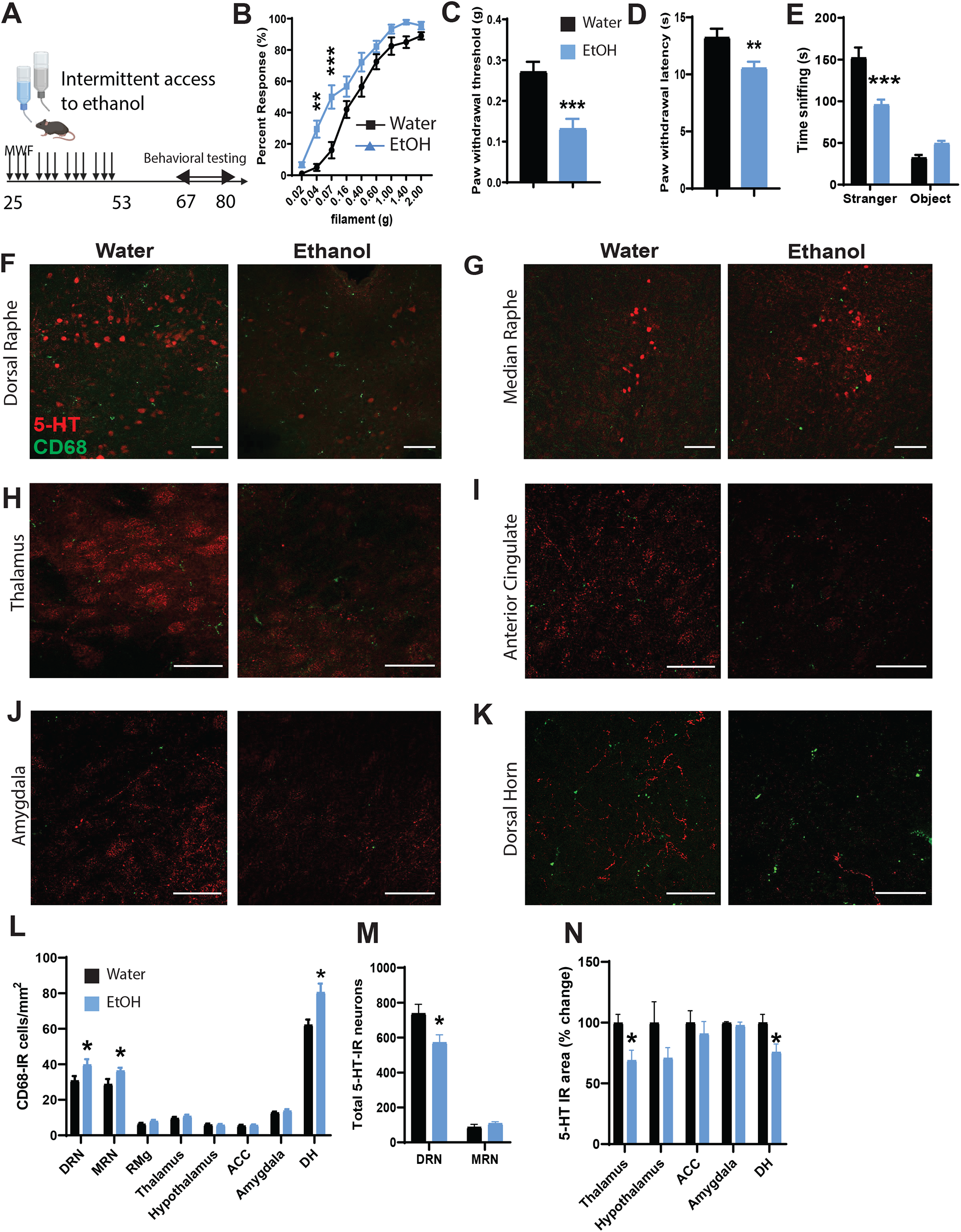
Persistent behavioral dysregulation and neuroinflammation in adult mice after adolescent alcohol exposure. (A) Experimental timeline for adolescent alcohol exposure. (B-C) Behavioral testing in the von Frey test, (D) Hargreaves test and (E) social interaction test in adult C57BL/6J mice after adolescent alcohol exposure. Confocal images of 5-HT and CD68 staining in the (F) DRN and (G) MRN and (H) thalamus, (I) ACC, (J) amygdala and (K) dorsal horn at 4 weeks after alcohol cessation. (L) Histogram of CD68+ cell counts, (M) 5-HT-IR cell counts in the DRN and MRN and (N) 5-HT-IR area in the thalamus, hypothalamus, ACC, amygdala, and dorsal horn. Scale bar = 100 µm in panels F-G and 50 µm in panels H-K. *p<0.05, **p<0.01, ***p<0.001.

### 3.2. AIE induces microglial activation across multiple brain regions involved in central pain processing

AIE increased the number of CD68-positive microglia in the DRN (t_23_=2.32, p<0.05) and MRN (t_20_=2.33 with Welch’s correction, p<0.05), suggesting that these microglia are in an activated state (Figures 1F, G and L). There was also a significant increase in the optical density of CD68 staining in the ACC (t_14_=2.61, p<0.05 with Welch’s correction) (Figure 1I, Supplementary Table 3). Surprisingly, there was no change in the number of CD68+ microglia or optical density in any other brain region examined (Supplementary Table 3).

P2Y12 is a purinergic receptor that is involved in platelet aggregation and is expressed at high levels in resting microglia. Once microglia are activated, P2Y12 expression decreases (Haynes et al., 2006). We found that adolescent ethanol reduced the number of P2Y12+ microglia in the thalamus (t_25_=3.01, p<0.01), amygdala (t_25_=2.20, p<0.05) and ACC (t_20_=2.46, p<0.05 with Welch’s correction), suggesting that microglia in these regions may also be in an activated state (Supplementary Figure 1 and Supplementary Table 3). P2Y12 is also transiently decreased in cortical microglia in response to an acute ethanol challenge in neonatal mice (Ahlers et al., 2015), so it is interesting that this P2Y12 reduction persists for several weeks after AIE cessation. There was also an increase in the area of P2Y12+ soma in the amygdala (t_25_=4.00, p<0.001) and the medullary raphe (t_22_=3.25, p<0.01 with Welch’s correction), which is consistent with an activated state.

Given the essential role of the spinal cord in nociception, we also examined CD68 and P2Y12 expression in the dorsal horn. There we found an increase in the number of CD68+ microglia in AIE mice (t_18_=3.28, p<0.01) (Figure 1K, L), but there was no change in the number of P2Y12+ microglia or soma size (Supplementary Figure 1, Supplementary Table 3).

The clinical and preclinical literature also suggests that chronic alcohol may reduce astrocyte density and nuclei size in the prefrontal and orbitofrontal cortex, ACC and hippocampus (Adermark and Bowers, 2016; Bull et al., 2015, 2014; Korbo, 1999; Miguel-Hidalgo et al., 2002). We did not observe a change in GFAP+ cell number or optical density in the DRN or MRN after AIE, but we did observe a reduction in cell body area in the DRN (t_18_=3.13, p<0.05 with Welch’s correction) (Supplementary Figure 2, Supplementary Table 3). We also observed an increase in the number of GFAP+ cells in the medullary raphe (t_17_=2.18, p<0.05) without any concomitant change in soma area.

### 3.3 AIE reduces the number of 5-HT-expressing neurons in the DRN as well as 5-HT immunoreactivity in the spinal cord and central pain processing regions

Next we examined 5-HT and Tph2 expression in the raphe nuclei after 4 weeks of withdrawal from AIE. There was a significant reduction in the total number of 5-HT-expressing neurons in the DRN (t_26_=2.48, p<0.05) (Figure 1M), but not the rate-limiting enzyme Tph2, suggesting that this may be driven by a reduction in tryptophan bioavailability or an increase in 5-HT metabolism. These results are in agreement with a previous study in rats exposed to adolescent ethanol gavage (Vetreno et al., 2017). Interestingly, there was no change in the number of 5-HT-IR or Tph2-IR neurons in the MRN and medullary raphe, although there was an increase in Tph2 optical density in the MRN (t_18_=3.18, p<0.01). We then decided to look downstream at nociceptive regions receiving 5-HT input from the DRN. 5-HT immunoreactive area was reduced in the thalamus (t_7_=2.92, p<0.05) and dorsal horn (t_18_=2.54, p<0.05) (Figure 1N), and there was also a reduction in 5-HT optical density in the thalamus (t_7_=2.98, p<0.05), ACC (t_7_=2.47=<0.05), and amygdala (t_7_=2.50, p<0.05).

### 3.4. Minocycline treatment alleviates pain and depressive-like behavior after AIE

Minocycline inhibits microglial activation *in vivo* and may have clinical utility in a variety of neurological conditions involving chronic inflammation. We found that AIE reduced mechanical pain thresholds in the Von Frey in vehicle-treated mice (Main effect of AIE: F_1,36_=5.50, p<0.05, Bonferroni post-test t_36_=2.73, p<0.05) which was abolished by chronic minocycline treatment after AIE cessation (t_36_=0.585, ns) (Figure 2A-C). Minocycline also normalized thermal sensitivity in the Hargreaves test (Main effect of alcohol F_1,56_=7.77, Alcohol x minocycline interaction F_1,56_=5.86, p<0.05) (Figure 2D). Bonferroni post-tests revealed an effect of AIE in vehicle-treated mice in the Hargreaves (t_56_=3.68, p<0.01) that was abolished in minocycline-treated mice (t_56_=0.26, ns).

**Figure 2:**
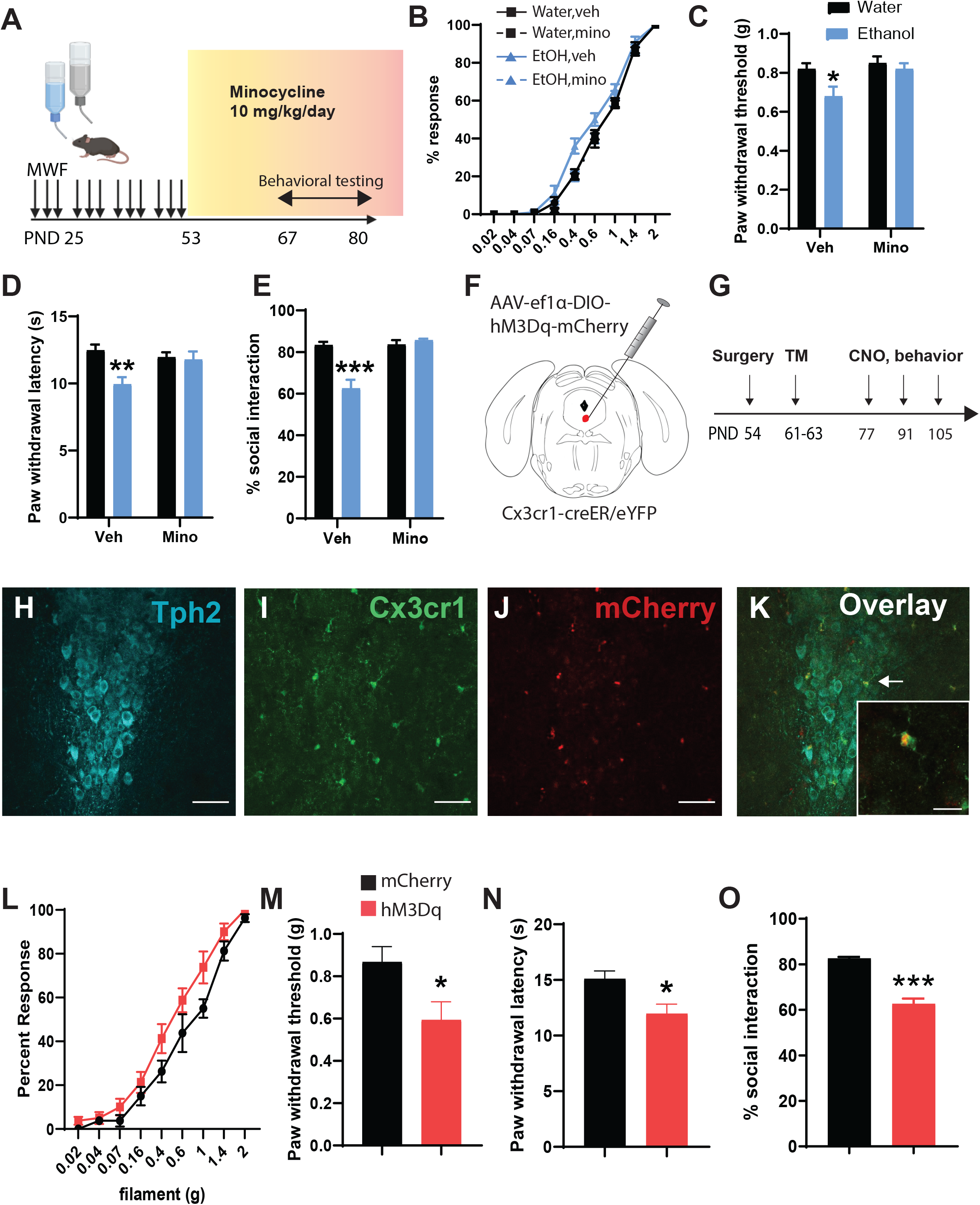
Minocycline rescues pain and depressive-like behaviors following adolescent alcohol exposure. (A) Experimental timeline for minocycline administration, adolescent alcohol exposure, and behavioral testing. (B-C) Percent response and histogram of paw withdrawal thresholds in the von Frey test, (D) Paw withdrawal latency in the Hargreaves test and (E) time spent in social interaction in minocycline and vehicle-treated C57BL/6J mice exposed to alcohol or water for 4 weeks in adolescence. (F) Stereotaxic injection of AAV expressing Gq-coupled DREADDs into the DRN of *Cx3cr1-creER* mice and (G) timeline of surgeries, tamoxifen and CNO injections, and behavior. (H-K) Confocal images of Tph2, GFP (expressed in Cx3cr1+ microglia), mCherry, and an overlay image showing colocalization of DREADDs with Cx3cr1+ microglia. Scale bar =50 µm (15 µm for inset). Behavior in the von Frey (L-M), Hargreaves (N) and (O) social interaction test after chemogenetic activation of DRN microglia. *p<0.05, **p<0.01, ***p<0.001.

Minocycline also rescued social interaction deficits in AIE-treated mice (F_1,36_=22.47, p<0.001 Ethanol x minocycline interaction). Pairwise comparisons revealed social deficits in ethanol vehicle-treated vs water vehicle-treated mice (t_36_=6.088, p<0.001) that were abolished by minocycline treatment (t_36_=0.6163, ns) (Figure 2E).

### 3.5. Chemogenetic activation of microglia in the DRN induces hyperalgesia and reduces social preference

We next asked whether direct activation of microglia in the DRN could phenocopy the effects of AIE. Male *Cx3cr1-creER* mice were microinjected with AAV-ef1α-DIO-hM3Dq-mCherry or a control virus (AAV-ef1α-DIO-mCherry) in the DRN at 8 weeks of age and then received i.p. injections of tamoxifen to enable Cre-mediated recombination and expression of the DREADDS in microglia (Figure 2F-K). Chemogenetic activation of microglia in ethanol-naïve *Cx3cr1-creER*::hM3Dq^DRN^ mice reduced mechanical and thermal pain thresholds in the Von Frey (t_14_=2.44, p<0.05) and Hargreaves test (t_14_=2.84, p<0.05) (Figure 2L-N). Social preference was also attenuated in these mice after microglial activation (t_14_=8.51, p<0.001) (Figure 2O).

## 4. Discussion

Our results indicate that AIE induces long-lasting adaptations in pain sensitivity and affective behavior that is accompanied by an increase in the number of activated (CD68-positive) microglia in the DRN and MRN. While CD68 expression was not significantly altered in other brain regions, we did observe a decrease in the number of P2Y12-expressing microglia in the thalamus, amygdala and ACC which have also been previously implicated in central pain processing. This was accompanied by an increase in the soma area of P2Y12+ microglia in the amygdala only. Surprisingly, there was no change in P2Y12+ microglia or soma area in the raphe nuclei, indicating that microglial activation states may differ between these brain regions. There was also a significant increase in CD68+ microglia in the dorsal horn of the spinal cord, suggesting that microglial activity in spinal and supraspinal pain circuits may contribute to persistent pain after adolescent alcohol exposure.

We then examined 5-HT expression in the DRN and MRN since activated microglia can release pro-inflammatory cytokines that have deleterious effects on 5-HT neurons (Hochstrasser et al., 2011b; Zhang et al., 2001). The number of 5-HT neurons was reduced in the DRN and accompanied by a reduction in 5-HT expression in the thalamus, ACC and amygdala. These regions were selected because they mediate descending modulation of pain and receive 5-HT input from the DRN (Ab Aziz and Ahmad, 2006; Chao et al., 2021; Duan et al., 2021; Gandhi et al., 2021; Gilpin et al., 2021; Heijmans et al., n.d.; Juarez-Salinas et al., 2019; Li et al., 2019; Price, 2000; Zargarani et al., 2021). There was also a decrease in 5-HT expression in the dorsal horn, which may have nociceptive or anti-nociceptive effects depending on which 5-HT receptors are prevalent (Sommer, 2010; Zhang et al., 2011). 5-HT neurons in the DRN were previously found to mediate anti-nociception (Dugué et al., 2014; Ruan et al., 1990), suggesting that depletion of 5-HT may contribute to increased pain thresholds in AIE mice. A previous study also found that 5-HT projections from the DRN to the parafascicular nucleus (Pf) of the thalamus inhibits nociceptive responses (Andersen and Dafny, 1983), so loss of 5-HT expression in the thalamus may also account for heightened pain sensitivity after AIE. Together, these studies suggest that the 5-HT DRN→Pf pathway may be a promising target for treating pain disorders with co-morbid alcohol dependence.

We also found that AIE-induced hyperalgesia could be reversed by systemic minocycline administration in adulthood, suggesting that targeting microglia may be an effective treatment strategy. This is supported by the finding that chemogenetic activation of microglia in the DRN increased mechanical and thermal sensitivity while reducing social preference. While previous studies have demonstrated that activating microglia in the entire brain and spinal cord can modulate nociception (Binning et al., 2020), our results demonstrate that regional activation of microglia in the DRN is sufficient to recapitulate the hyperalgesia observed following AIE. Chemogenetic activation of microglia also stimulates the release of pro-inflammatory cytokines which can have direct effects on 5-HT neuronal activity (Brambilla et al., 2007; Manfridi et al., 2003). In another study, overexpression of IL-1β in the DRN resulted in manic-like behavior characterized by reduced fearfulness and increased stress-induced locomotor activity (Howerton et al., 2014). While we did not test mania-like behaviors directly or measure IL-1β levels in the DRN, previous studies have shown that IL-1β is elevated in other brain regions after chronic ethanol exposure (Bajo et al., 2015). Overall, our studies support the conclusion that adolescent alcohol drinking can increase the risk of chronic pain by activating microglia in the DRN and reducing 5-HT neurotransmission. Pharmacological interventions aimed at reducing neuroinflammation or increasing 5-HTergic tone may have therapeutic potential in the treatment of pain disorders in patients with a history of adolescent alcohol use.

## Supporting information

Supplemental Methods and Results

## Acknowledgements

This work was funded through NIH grants R00 AA024215, R01 AA028931 and BBRF grant #27530 to C.M. K.K. was supported by T32 DK112751.

