## Supplemental Methods and Results for "Adolescent ethanol drinking promotes hyperalgesia, neuroinflammation and serotonergic deficits in mice that persist into adulthood"

**Supplementary Material**

### Inventory of Supplemental Information

#### Supplementary Methods

Intermittent access to ethanol

Open Field

Elevated Plus Maze

Forced Swim

Image Analysis

#### Supplemental data

Supplementary Table 1: Summary of results from the elevated plus maze, open field and forced swim test after adolescent alcohol

Supplementary Table 2: antibodies used in IHC

Supplementary Table 3: Summary of histology results after adolescent alcohol

Supplementary Figure 1: Representative images of P2Y12 expression in the brain and spinal cord following adolescent IA

Supplementary Figure 2: Representative images of GFAP expression in the raphe nuclei following adolescent IA

**Supplementary Methods**

**Intermittent access to ethanol**

On the first four days of the experiment, ethanol content was 3%, 6%, 10%, and 20% w/v with 5% sucrose. On day 5, animals received 20% EtOH w/v & 2% sucrose. Thereafter, ethanol content was 20% w/v.

**Behavior tests:**

Following intermittent access to ethanol, animals entered into a chronic withdrawal period. Behavior testing began ~3 weeks into chronic withdrawal.

**Open Field**

Mice were placed in the corner of a 25 x 25 x 25 cm plexiglass container as described previously and allowed to freely explore the chamber for 10 minutes (Marcinkiewcz et al., 2019, 2016, 2014). The time spent in the center of the container as well as total distance traveled was recorded by an overhead camera integrated with Ethovision software. The center of the open field was defined as the central 10% of the arena.

**Elevated Plus Maze**

Mice were placed in the center of an elevated plus maze and allowed to freely explore the arena for 5 minutes. The lux in the open arms were roughly ~20 lux. Time spent in the open arms and the number of transitions between closed and open arms was recorded by an overhead camera and quantified with Ethovision software.

**Forced Swim**

Mice were gently placed in a tall cylinder (32 cm height x 20 cm diameter) filled with tap water ($24^0$C) to a height of 25cm, for 6 minutes. After the 6 minutes, animals were removed from the test cylinder and placed in a clean cage under a heating lamp for 5-minutes to warm them up and allow them to dry off. Behaviors were recorded from a side-view camera and analyzed with Ethovision software. Latency to immobility, as well as number of immobile bouts + total immobile time during minutes 0-2 (‘pretest’) and min 2-6 (‘test’) were evaluated.

**Image Analysis:**

Images were imported to ImageJ and analyzed by trained researchers blind to experimental conditions for cell counts, cell body size, and optical density.

The intensity of serotonergic cells and glia were estimated via optical density measurement in ImageJ. Briefly, images were converted to 8-bit gray scale images and background was corrected. Mean gray values were obtained from regions of Interest (ROIs) that were overlaid on the grayscale images, with an effort to avoid obvious artifacts.  The same ROIs were overlaid on z-stacks to quantify the number of 5HT, tph2, GFAP, CD68, and P2Y12 immunoreactive cells. Morphological analysis was performed on GFAP and P2Y12 positive cells; observers blind to the experimental groups traced outlines of the glial cells, making sure to avoid dendritic processes.

| Assay | Measure | *p*-value |
|---|---|---|
| Open Field | Distance moved (cm) | ns |
| | Time in the center (s) | **.0200 (*)** |
| Elevated Plus Maze | Distance moved (cm) | ns |
| | Time spent in the open arms (s) | ns |
| | Time spent in the open arms (%) | ns |
| | Probability to enter the open arms | ns |
| Forced Swim test | Latency to inactivity (s) | ns |
| | Pretest: inactive frequency | ns |
| | Pretest: inactive time (s) | ns |
| | Pretest: average length of inactive bouts (s) | ns |
| | Test: inactive frequency | **.0422 (*)** |
| | Test: inactive time (s) | Ns |
| | Test: average length of inactive bouts (s) | **.0417 (*)** |

**Supplementary Table 1: Summary of p values for statistical tests comparing data from adolescent alcohol and control mice in the open field, EPM, and forced swim tests.** *P*-values less than .05 have been bolded. * denotes $p < .05$.

| Stain | Antibody | Concentration |
|---|---|---|
| 5HT-CD68-P2Y12 | Goat anti-5HT<br>(Immunostar; catalog # 20079) | 1:4000 |
| | Rat anti-CD68<br>(Abcam; catalog # ab53444) | 1:500 |
| | Rabbit anti-P2Y12<br>(Anaspec; catalog # A5-55043A) | 1:2000 |
| | Cy3 Donkey anti-Goat<br>(Jackson ImmunoResearch; catalog # 705-165-147) | 1:500 |
| | Alexa Fluor 488 Donkey anti-rat<br>(Jackson ImmunoResearch; catalog # 712-545-150) | 1:500 |
| | Alexa Fluor 405 Donkey anti-rabbit<br>(Jackson ImmunoResearch; catalog # 711-475-152) | 1:500 |
| 5HT-tph2-GFAP | Goat anti-5HT<br>(Immunostar; catalog # 20079) | 1:2000 |
| | Rabbit anti-tph2<br>(Novus; catalog # NB100-74555) | 1:1000 |
| | Chicken anti-GFAP<br>(Abcam; catalog # ab4674) | 1:6000 |
| | Cy3 Donkey anti-Goat<br>(Jackson ImmunoResearch; catalog # 705-165-147) | 1:500 |
| | Alexa Fluor 488 Donkey anti-chicken<br>(Jackson ImmunoResearch; catalog # 703-545-155) | 1:500 |
| | Alexa Fluor 405 Donkey anti-rabbit<br>(Jackson ImmunoResearch; catalog # 711-475-152) | 1:500 |

**Supplementary Table 2: Antibodies used in IHC**

| | Tph2 | | GFAP | | | CD68 | | P2Y12 | |
|---|---|---|---|---|---|---|---|---|---|
| | Number of cells | Optical density | Number of cells | Cell body size | Optical density | Number of cells | Optical density | Number of cells | Cell body size |
| *Dorsal Raphe Nucleus* | ns | ns | ns | **.0101 (*)** | ns | **.0299 (*)** | ns | ns | ns |
| *Median Raphe Nucleus* | ns | **.0051 (**)** | ns | ns | ns | **.0303 (*)** | ns | ns | ns |
| *Medullary Raphe* | ns | ns | **.0436 (*)** | ns | ns | ns | ns | ns | **.0037 (**)** |
| *Thalamus* | — | — | — | — | — | ns | ns | **.0059 (**)** | ns |
| *Amygdala* | — | — | — | — | — | ns | ns | **.0376 (*)** | **.0005 (***)** |
| *Hypothalamus* | — | — | — | — | — | ns | ns | ns | ns |
| *Anterior Cingulate* | — | — | — | — | — | ns | **.0208 (*)** | **.023 (*)** | ns |
| *Dorsal Horn* | — | — | — | — | — | **.0042 (**)** | ns | ns | ns |

**Supplementary Table 3: Summary of p values for statistical tests comparing Tph2, GFAP, CD68 and P2Y12 expression in adolescent alcohol mice relative to control mice.** Values where $p < .05$ have been bolded. * denotes $p < .05$, ** denotes $p < .01$, *** denotes $p < .001$

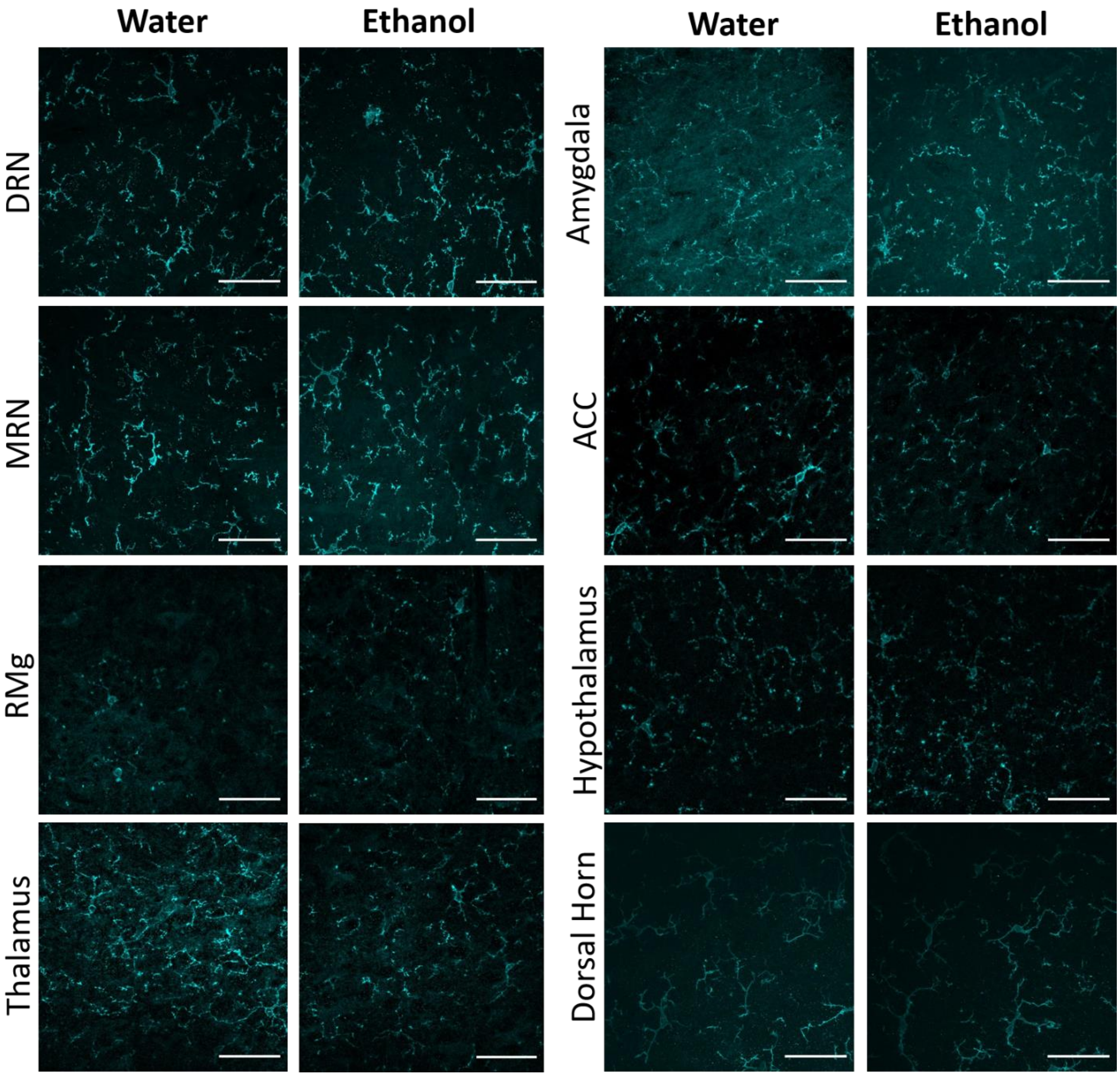


**Supplementary Figure 1: Representative images of P2Y12 in the DRN downstream regions following adolescent IA.** We observed a significantly reduced number of P2Y12 positive cells in the Thalamus, Amygdala, and Anterior Cingulate following adolescent IA. We also observed greater cell body area of P2Y12 positive cells in the Amygdala. Scale bar: 50µm.

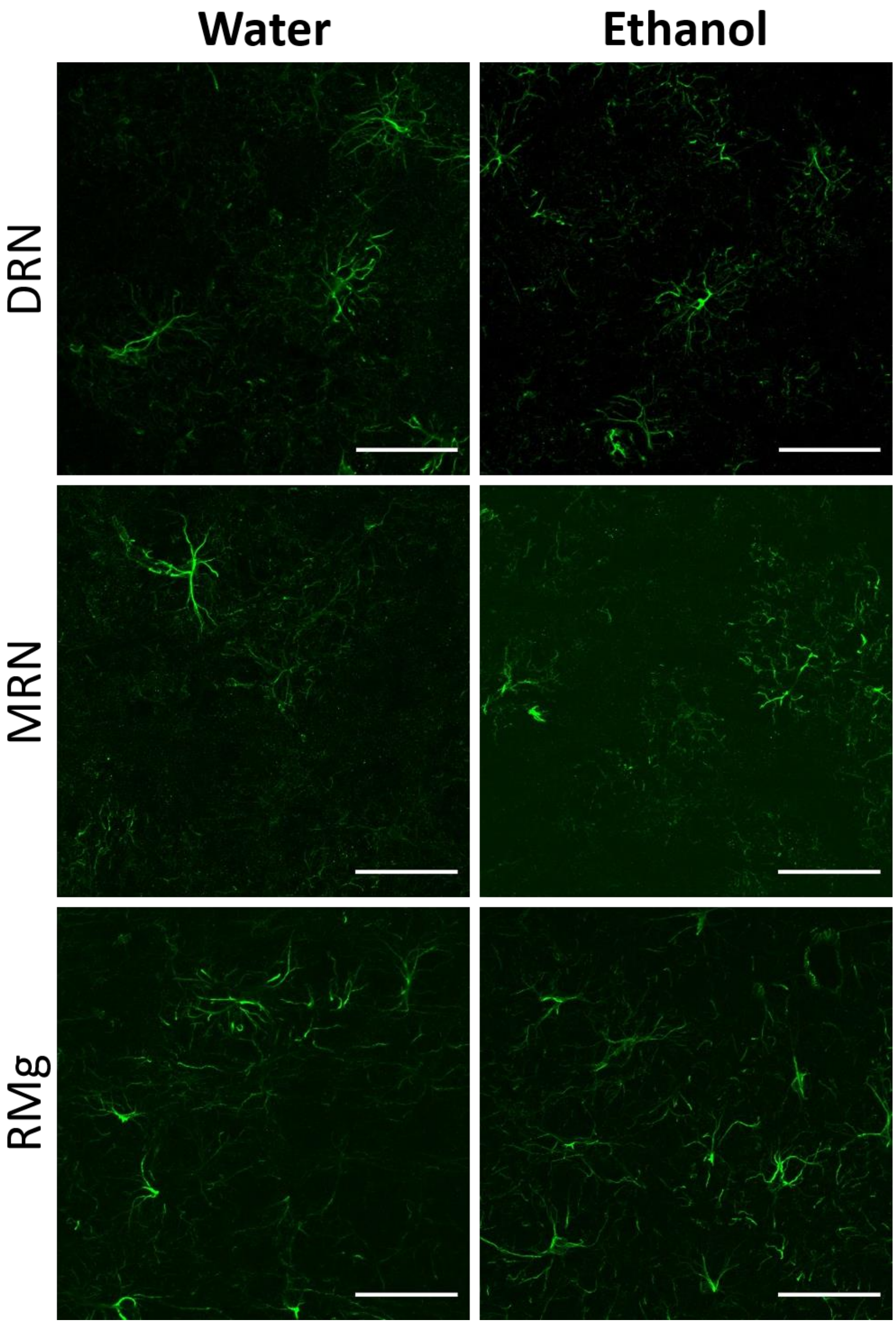


**Supplementary Figure 2: Representative images of GFAP in the raphe nuclei following adolescent IA.** We observed a statistically significant increase in cell body size in the DRN following adolescent alcohol, and an increase in the number of GFAP positive cells following adolescent alcohol exposure. Scale bar: 50µm.
